# hyve, a compositional visualisation engine for brain imaging data

**DOI:** 10.1101/2024.04.18.590179

**Authors:** Rastko Ciric, Anna Xu, Russell A. Poldrack

## Abstract

Visualisations facilitate the interpretation of geometrically structured data and results. However, heterogeneous geometries—such as volumes, surfaces, and networks—have traditionally mandated different software approaches. We introduce `hyve`, a Python library that uses a compositional functional framework to enable parametric implementation of custom visualisations for different brain geometries. Under this framework, users compose a reusable visualisation protocol from *geometric* primitives for representing data geometries, *input* primitives for common data formats and research objectives, and *output* primitives for producing interactive displays or configurable snapshots. `hyve` also writes documentation for user-constructed protocols, automates serial production of multiple visualisations, and includes an API for semantically organising an editable multi-panel figure. Through the seamless composition of input, output, and geometric primitives, `hyve` supports creating visualisations for a range of neuroimaging research objectives.

## 1 Introduction

By associating data with geometric structures, visualisation provides a gateway to understanding and communicating complex information. In neuroimaging, embedding data in brain geometries improves the tractability of localising and interpreting high-dimensional observations and results. Visualisation software (e.g., Claudi et al., 2021; Combrisson et al., 2019; Xia et al., 2013) is thus critically important in the discipline of brain mapping (Chopra et al., 2023).

However, the geometries that structure neuroimaging datasets can be highly heterogeneous. Magnetic resonance data, for example, are often reconstructed as image intensity values in a regularly sampled three-dimensional Euclidean volume. By contrast, the convoluted sheet of the mammalian cerebral cortex can be modeled as a two-dimensional Riemannian manifold, which software suites approximate as a polygon mesh (or surface; Greve et al., 2014; Pang et al., 2023). Maps of brain connectivity have the intrinsic topology of a sparse or dense graph, with vertices that can be embedded either in physical coordinates or algorithmically in a low-dimensional space so as to preserve proximity of strongly connected vertices (Fanton and Thompson, 2023; Xia et al., 2013). Measures of functional activity in the brain further extend these geometries in the dimension of time.

Existing software has often addressed this heterogeneity by implementing separate plotting routines for different geometries. Here, we introduce the software library `hyve` (the `hypercoil` visualisation engine) to implement an alternative approach that leverages compositional functional programming. In our compositional paradigm, users construct a new visualisation protocol by composing an abstract base plotting routine with a chain of functional atoms called *primitives*. Each functional primitive imbues the base routine with a distinct functionality, forming the basis of a modular, flexible, and extensible system for building reusable plotting protocols.

`hyve` includes *input primitives* to facilitate the loading and transformation of common data types, as well as *output primitives* that determine the form in which the visualised scene is ultimately represented—for instance, an interactive window, PNG snapshots of fixed views, or an editable multi-panel SVG figure. This collection of primitives was designed with the intention to support a diverse range of data geometries and scientific objectives, initially motivated by our work on a differentiable programming pipeline for neuroimaging data (Ciric, Thomas, et al., 2022).

In this article, we begin by introducing the overall design principles and organisational schema of `hyve`. Next, we discuss the software from the perspective of a typical user, before proceeding to technical implementation details. From the perspective of both *use* and *implementation*, we review currently supported geometries. For each geometry, we highlight some input primitives that can be used to meet common scientific objectives. Finally, we provide an overview of output primitives, including the `hyve` figure builder.

## 2 Software overview for users

### 2.1 Design principles

`hyve` is an open-source Python library for creating visualisations of neuroimaging data. Python is the *de facto* high-level language of choice for non-proprietary software projects in neuroimaging, with a mature software ecosystem that has wide currency in the community (e.g., Abraham et al., 2014; Brett et al., 2023; Gorgolewski, 2011). Python also has a number of flexible libraries that provide intuitive, high-level bindings to software systems for graphics and rendering (e.g., Musy et al., 2023; Ramachandran and Varoquaux, 2011; Sullivan and Kaszynski, 2019).

The versatility of the `hyve` visualisation system comes from compositional functional programming. Under this programming paradigm, complex visualisation protocols are constructed by composing atomic functions, called primitives, in a combinatorial manner. The collection of available primitives together form a high-level grammar for plot definition (Wilkinson, 2005), for which `hyve` is the interpreter; this model of graphics generation takes loose inspiration from the *ggplot* library within the R software ecosystem. Composition is performed under `hyve.plotdef`, the main function of `hyve`’s user interface (Figure 1).

**Figure 1:**
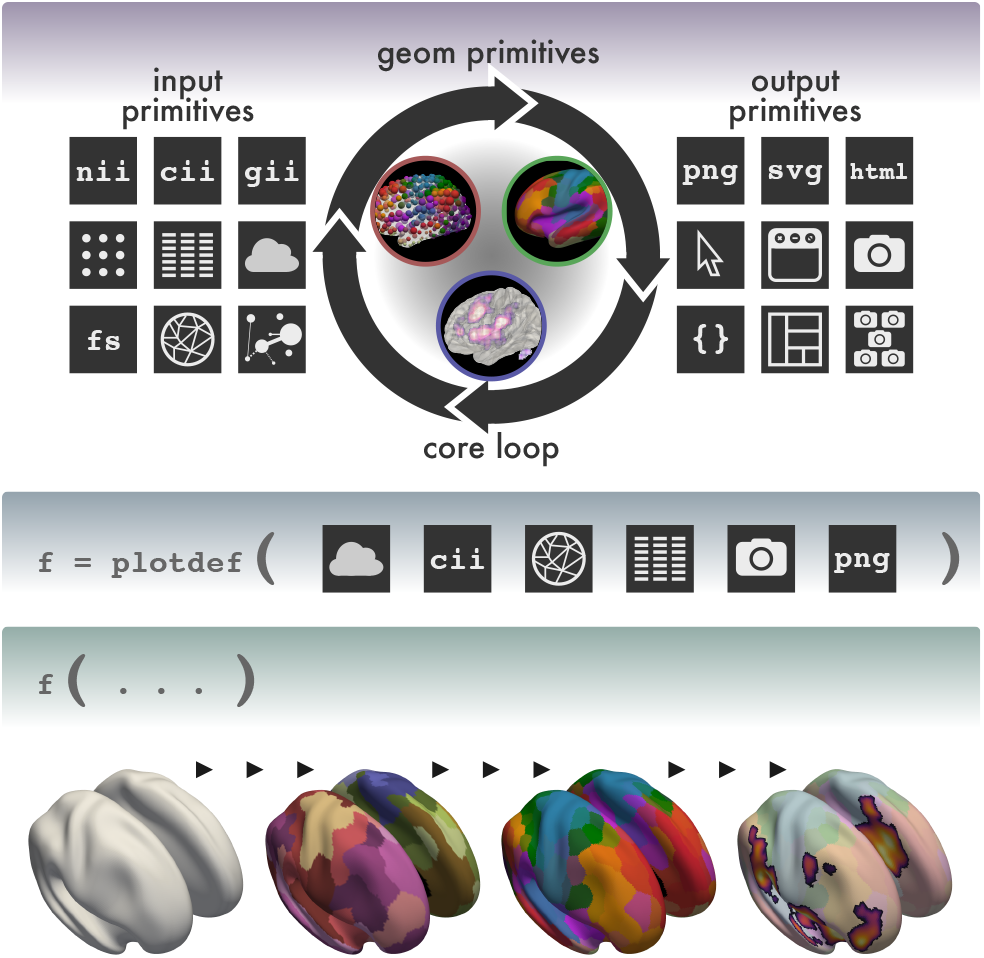
Overall framework of `hyve`. *Top*, `hyve` is a visualisation engine for neuroimaging data. `hyve`’s base plotter is composed with geometric primitives that interpret neuroimaging datasets structured by geometries such as surfaces, volumes, and networks. The plotter is wrapped in a core loop that returns scenes and associated metadata by mapping argument values over multiple plotter calls. Input and output primitives change the functionality of the core loop. *Centre*, the principal function in the `hyve` API is `plotdef`. `plotdef` takes as arguments a chain of transforms, each of which composes a particular primitive into the plotter. *Bottom*, `plotdef` returns a new function that implements a particular visualisation protocol. The function is reusable and automatically documented. As additional primitives are composed into the function, expressive visualisation styles become available.

#### 2.1.1 Basic usage

Not yet having introduced any particular functional primitives, we provide a preview of `hyve` usage. First, `hyve.plotdef` is invoked, with a sequence of positional parameters corresponding to the input and output primitives to be composed—in this example, to define a protocol for interactively visualising cortical parcellations. As shown below, some primitives must be parameterised before they are composed into the visualisation loop. `plotdef` returns a new visualisation protocol—a *function* that we have called `plot_f`.

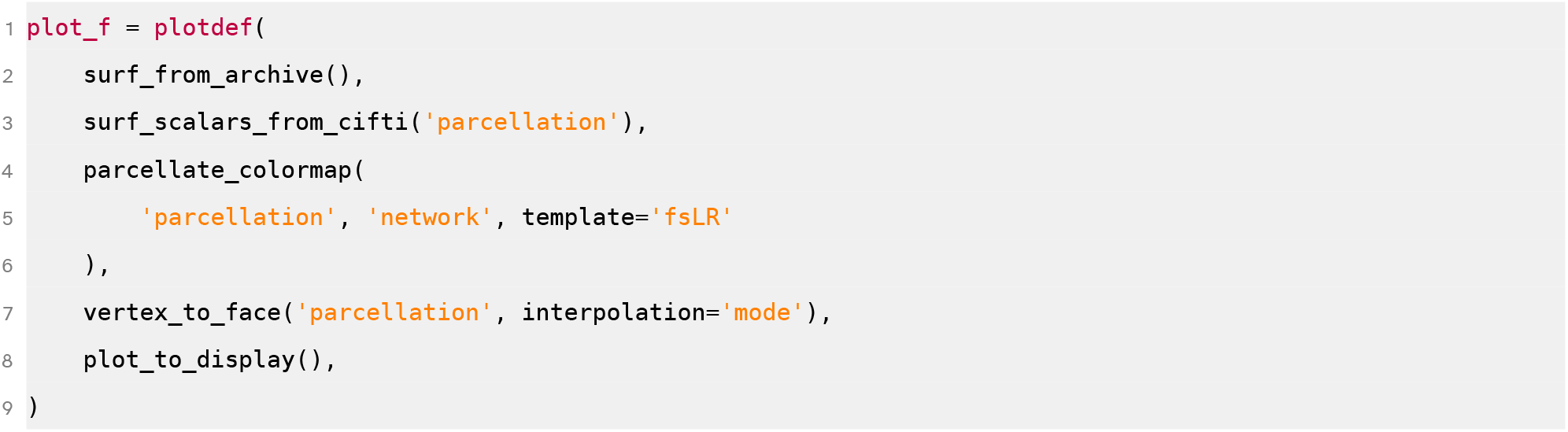

This protocol can then be invoked and re-invoked with new arguments like any Python function, for instance to return a different view of a different parcellation:

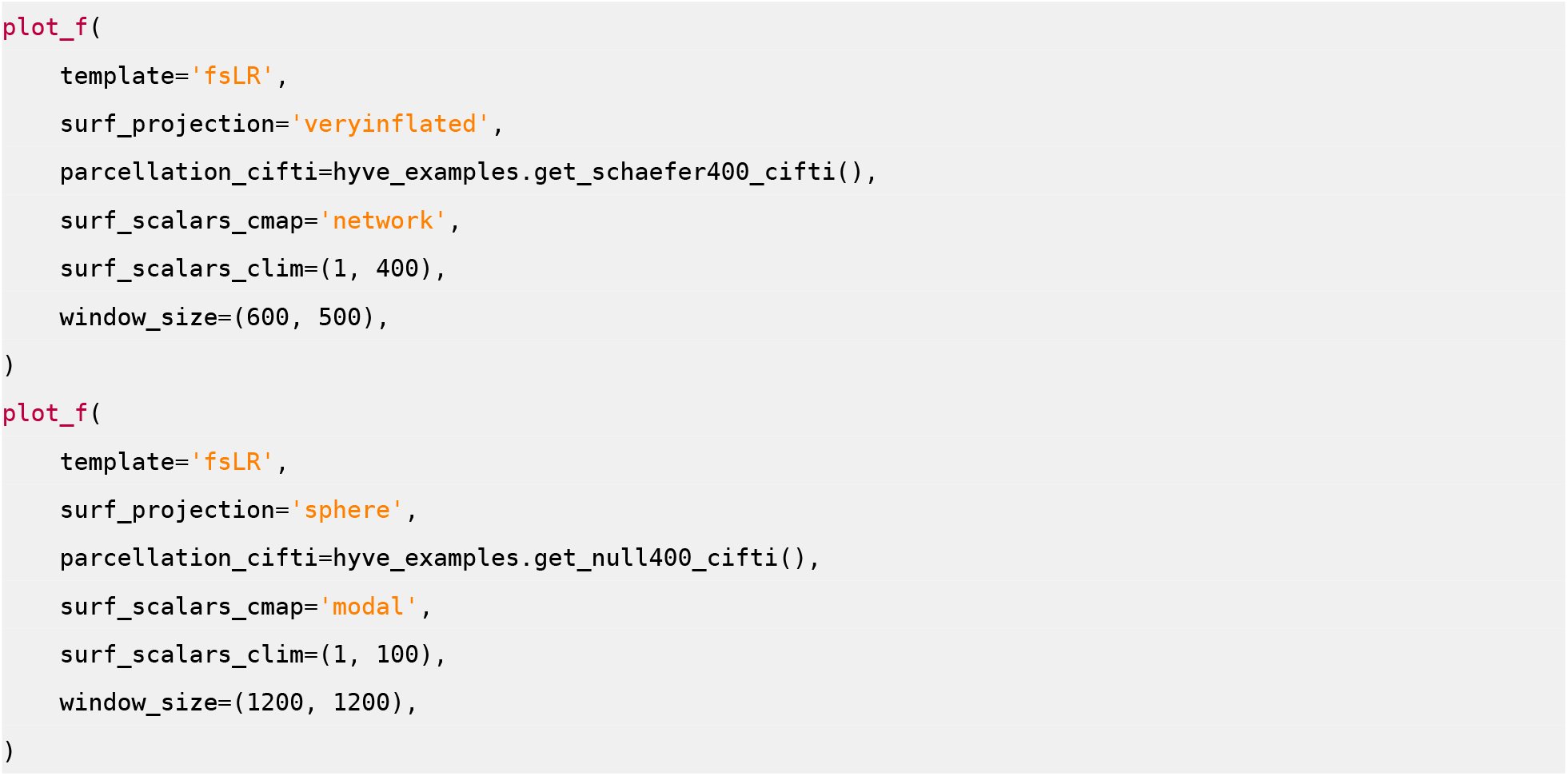

Moreover, `plot_f` has documentation and type hints that can be viewed using `help(plot_f)`, and it supports parameter auto-completion in interactive Python sessions. Note that, under the functional programming paradigm, both the arguments (primitives) and the return value (visualisation protocol) of `plotdef` are functions.

#### 2.1.2 How it works

All `hyve` visualisation protocols ultimately call a core plotting routine. This core routine is a visualisation *loop*: each invocation of a plotting protocol has the flexibility to render multiple scenes. This design enables, for instance, a single call to produce all six orthographic views on scenes that show both cerebral hemispheres together and each hemisphere in isolation, and afterward to collate all of these products in a single “panoptic” figure.

When a protocol receives a sequence (e.g., a Python tuple) as an argument, it will map the plotter over each value of that argument, with the capacity for customisable combinatorial mapping (see Section 3.1.2 and Figure 8 for details). To return to the previous example, providing the argument `hemisphere=(‘left’, ‘right’, ‘both’)` will repeat the visualisation procedure three times, once selecting only left hemisphere data, once selecting only right hemisphere data, and once showing both hemispheres.

`hyve.plotdef` composes a sequence of *input* and *output primitives* into the core visualisation loop. Input primitives transform the original loop, which operates on `hyve`-internal geometry containers, into a new function that operates on input types that are commonly used in neuroimaging research, or else they add functionality that fulfils some research objective (like investigating parcellations or highlighting brain subnetworks—discussed later). Input primitives (e.g., `surf_scalars_from_cifti`) are available for reading data from `numpy` arrays or images in NIfTI, CIfTI, GIfTI, or FreeSurfer formats into a surface geometry container. Correspondingly, output primitives (e.g., `plot_to_display`) transform the original loop, which returns a basic plotter object for each scene, into a new function that serialises scene representations into interpretable output formats. These formats include interactive display windows and HTMLs, static snapshots of selected views on a scene, or editable SVG figures with different scenes arranged in panels.

### 2.2 Data geometries and input primitives

`hyve` can visualise data geometries that commonly occur in neuroimaging: surfaces, volumes, and brain networks. Across these geometries, a dataset’s values can be mapped to different visual properties of its structuring geometry (or “aesthetics”; Wilkinson, 2005). Any visualisation protocol defined by `hyve.plotdef` automatically acquires parameters for configuring these properties, which will likely be familiar for users of high-level visualisation frameworks. Examples include:

- `color`: sets the geometric representation uniformly to the specified colour;
- `cmap`: maps the values of a scalar-valued dataset into a path in RGBA space defined in the specified colour map;
- `clim`: configures the limits of the dynamic range of the colour map;
- `cmap_negative` and `clim_negative`: maps negative values of the dataset using a separate colour map;
- `radius`: sets the radius of a mesh object (e.g., spheres and cylinders in network scenes), either uniformly or according to a specified scalar-valued dataset;
- `rlim`: configures the limits of the dynamic range of the radius;
- `alpha`: sets the opacity of the visualised object, either uniformly or according to a specified scalar-valued dataset (option not available for all geometries);
- `below_color`: sets mesh elements whose data values are below the minimum of the dynamic range uniformly to the specified colour.

By convention, each aesthetic parameter is prefixed with a designation for the geometry that it configures (e.g., `surf_scalars_cmap`, `points_scalars_cmap`, `node_cmap`, `edge_cmap`).

#### Geometric scalar datasets

Scalar datasets defined over geometric structures are abundant in neuroimaging: they include *inter alia* statistical maps, hard parcellations, and probability priors. Vector datasets, like signal decompositions and time series, can also be sampled to produce a sequence of scalar-valued slices. Scalar data are often defined either over voxels in a Euclidean volume or vertices on a surface mesh. Visualisation of common scalar data types is discussed below.

#### Surface scalars

Surface scalar datasets (Figure 2) require an associated mesh surface to structure them; `hyve` provides input primitives for either fetching this surface as a standard template from a remote archive (using `templateflow` or a `neuromaps` fork; Ciric, Thompson, et al., 2022; Markello et al., 2022) or providing it explicitly in GIfTI or FreeSurfer format (Fischl, 2012). Scalar datasets are then associated with surface vertices using primitives that support input from GIfTI, CIfTI, and NIfTI formats. Datasets in NIfTI format must be in approximate register with the `MNI152NLin2009cAsym` space; they are resampled to the surface using a `neuromaps`-based transformation (Markello et al., 2022).

**Figure 2:**
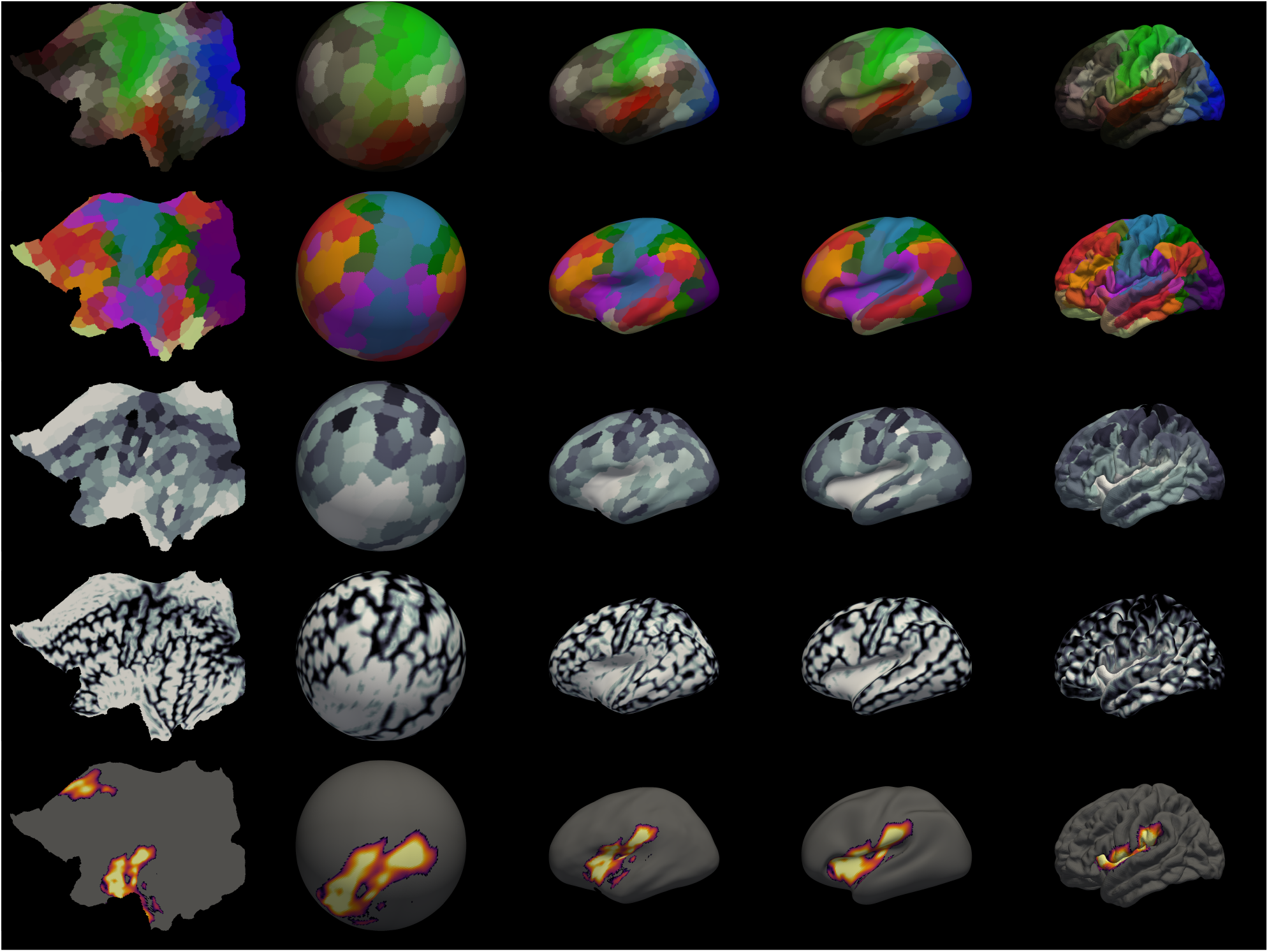
Selected static visualisation configurations of the geometric primitive for surface scalars. Above are plotted 25 simple, static `hyve` visualisations of surface scalar data on the left cortical hemisphere. Each column corresponds to a different surface projection—flat, spherical, two inflated projections, and pial. Each row corresponds to a different configuration of inputs and input primitives. The first three rows plot the 400-region gwMRF parcellation from Schaefer et al., 2017. The first shows the `modal` colour map, while the second shows the `network` colour map; these are obtained from surface-bound RGBA datasets included in `hyve` using the `parcellate_colormap` input primitive. The third row colours each parcel according to the mean value of a grey matter density dataset within that parcel; this behaviour is implemented in the `parcellate_surf_scalars` input primitive. The fourth row shows the original vertex-valued grey matter density dataset (Ciric, Thompson, et al., 2022) for comparison. The fifth row shows a statistical map from a meta-analysis of pain (Xu et al., 2020), resampled from a volumetric MNI space using the `surf_scalars_from_nifti` input primitive.

One common type of surface dataset is the cortical parcellation, and `hyve` provides additional input primitives that facilitate working with parcellations. `parcellate_colormap` creates a custom colour map for any parcellation dataset by parcellating a RGBA dataset inspired by prior literature. Two source datasets are available in `fsLR` and `fsaverage` space:

- The `network` RGBA dataset assigns each vertex a colour inspired by the 7-network parcellation from Yeo et al., 2011.
- The `modal` RGBA dataset assigns each vertex a colour inspired by the modal annotation from Glasser et al., 2016 (visual=blue, auditory=red, somatomotor=green).

`scatter_into_parcels` transforms the core loop to visualise an array defined over parcels instead of one defined over vertices, and then scatters each parcel’s data value across all vertices assigned to that parcel. `parcellate_surf_scalars`, by contrast, maps a dataset that is defined over the vertices of a surface into parcels.

#### Point cloud scalars

Volumetric datasets can either be resampled onto the surface (and thus interpreted as surface scalars, at the cost of subcortical information) or represented as *point cloud scalars*. Point cloud scalars are read from a NIfTI source file using an input primitive (`point_scalars_from_nifti`). They can be placed in a scene together with a low-opacity surface in order to produce “glass brain” visualisations of a dataset (Figure 3, *top*), which can provide intuition into the 3-dimensional shape and extent of brain maps, complementary to the 2-dimensional information presented in single-slice visualisations.

**Figure 3:**
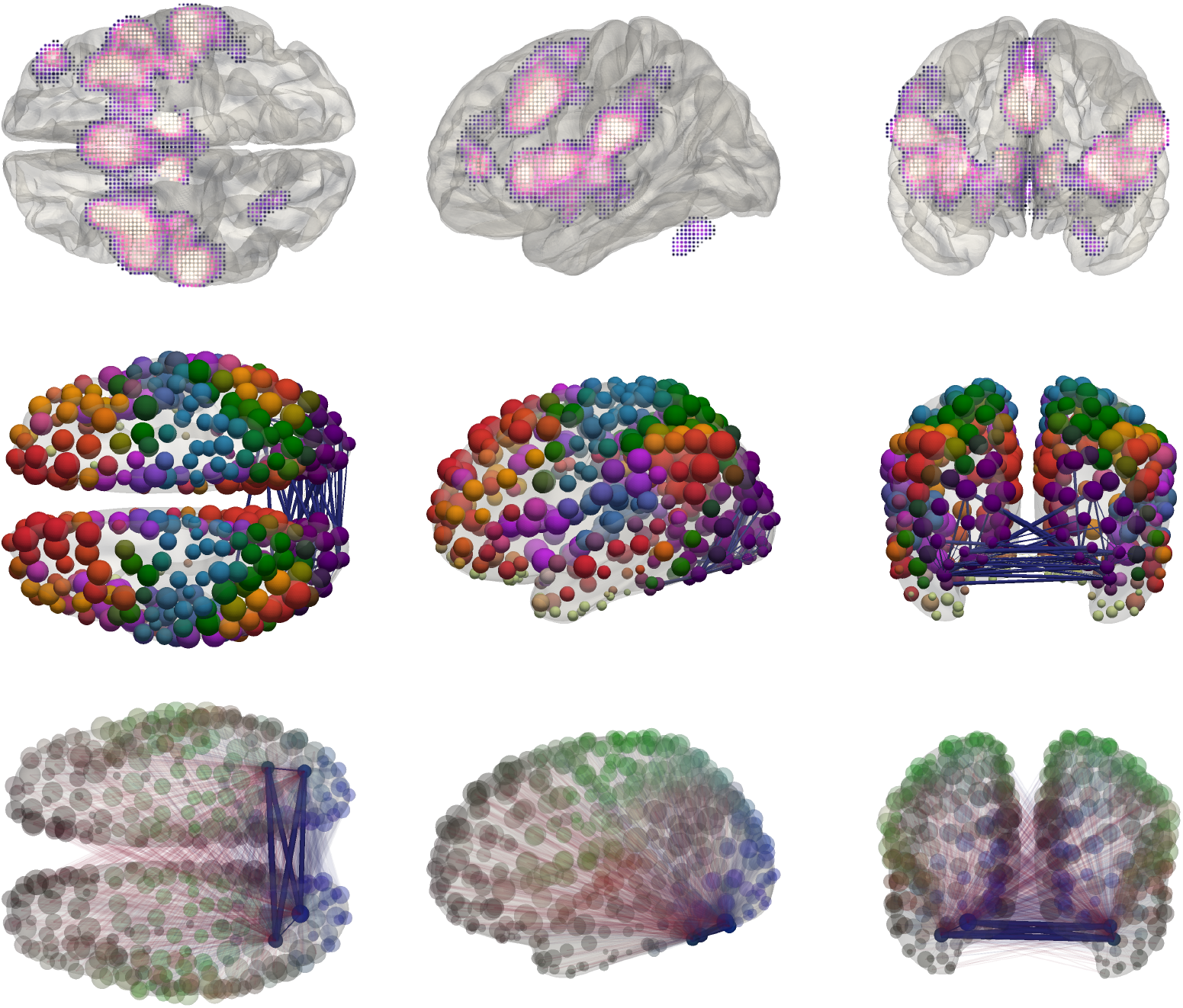
Selected static visualisation configurations of the geometric primitives for point scalars and brain networks. The top row shows “glass brain” plots of a statistical map from a meta-analysis of pain (Xu et al., 2020), loaded using the `points_scalars_from_nifti` input primitive. The second and third rows show a visual subnetwork from a synthetic connectome (Laumann et al., 2016) over the 400-region Schaefer parcellation (Schaefer et al., 2017). In the middle row, nodes are coloured using the `network` colour map and sized according to overall degree using the `add_node_variable` input primitive. A subset of visual network edges, sized by strength, is selected and displayed using the `add_edge_variable` input primitive. In the bottom row, all edges incident on a subset of four visual nodes are plotted, with edges scaled by strength and coloured by sign. The four nodes and their internal connections are focused by modulating the `alpha` property.

#### Brain networks

The brain network data that frequently occur in connectomics present a persistent challenge for visual representation. Following previous work (e.g., Fanton and Thompson, 2023; Xia et al., 2013), `hyve` uses a ball-and-stick scheme wherein nodes are represented as sphere meshes and edges as cylinder meshes (Figure 3, *middle and bottom*). `hyve` includes primitives for constructing node- and edge-valued variables from `numpy` arrays and `pandas.DataFrame`s, including support for thresholding operations and highlighting subsets of edges and nodes. Aesthetic properties of the ball-and-stick representation, such as the width and colour of elements, can be mapped to node degree, edge weight, or any variable defined over the network’s edges and nodes. Technical details about data representation are provided in Section 3.2.2, together with an overview of available filters and primitives.

#### Overlays

It is also possible to visualise multiple datasets as layers over a single geometry, for instance to investigate the spatial overlap between different brain maps (Figure 4). For each geometry, `hyve` includes an `add_*_overlay` primitive. `add_*_overlay` is parameterised by (i) a unique overlay name and (ii) a chain of input primitives that are applied specifically in the context of the overlay. For example, the below plot definition situates two overlays called `parcellation` and `statmap` on a surface geometry, and applies a different set of transforms to each.

**Figure 4:**
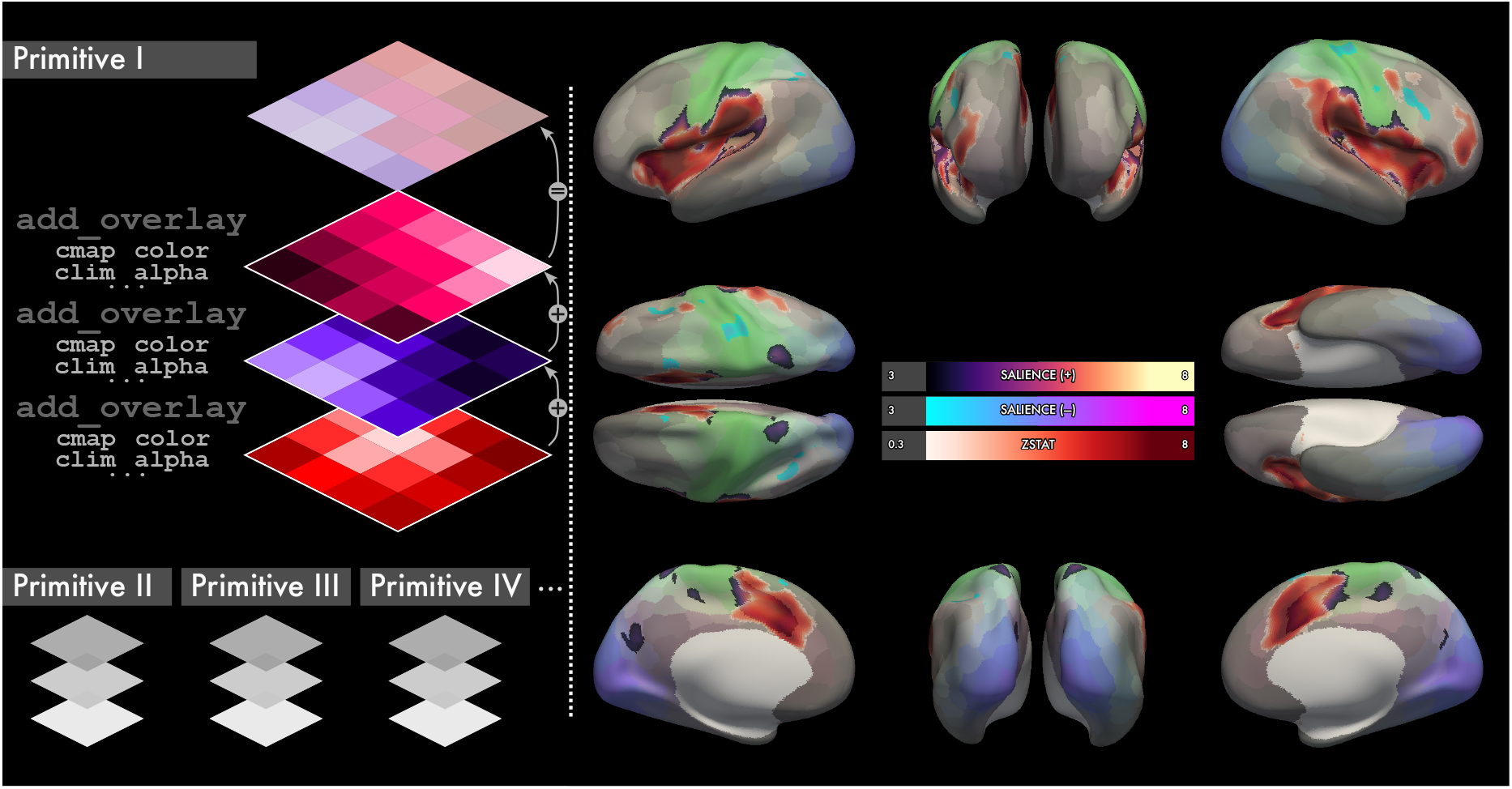
The `hyve` overlay system. For each geometric primitive, `hyve` supports adding multiple overlays, each of which transforms the function returned by `hyve.plotdef` to accept arguments for configuring the overlay’s visual properties. The data in overlays are blended together into a RGBA tensor using the *source-over* operation. The panoptic spread at right demonstrates the system in action: a low-opacity view on the 400-region Schaefer atlas is plotted as the base layer (Schaefer et al., 2017). The next layer contains a thresh-olded map of an independent component corresponding to the salience network. Finally, the top layer contains a thresholded z-statistic map of pain-related brain regions from the meta-analysis of Xu et al., 2020.

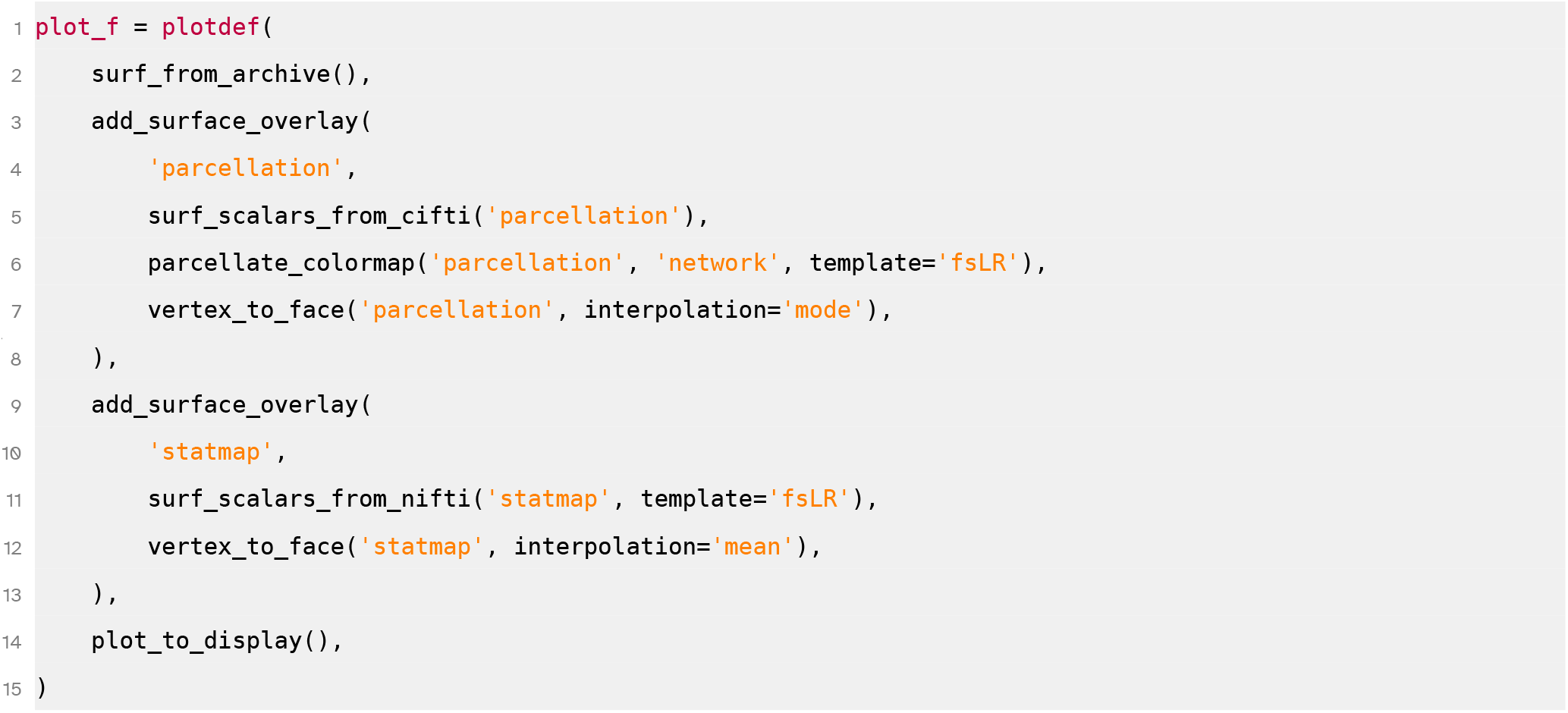

The specified overlay name then determines the names of new aesthetic arguments added to the visualisation protocol:

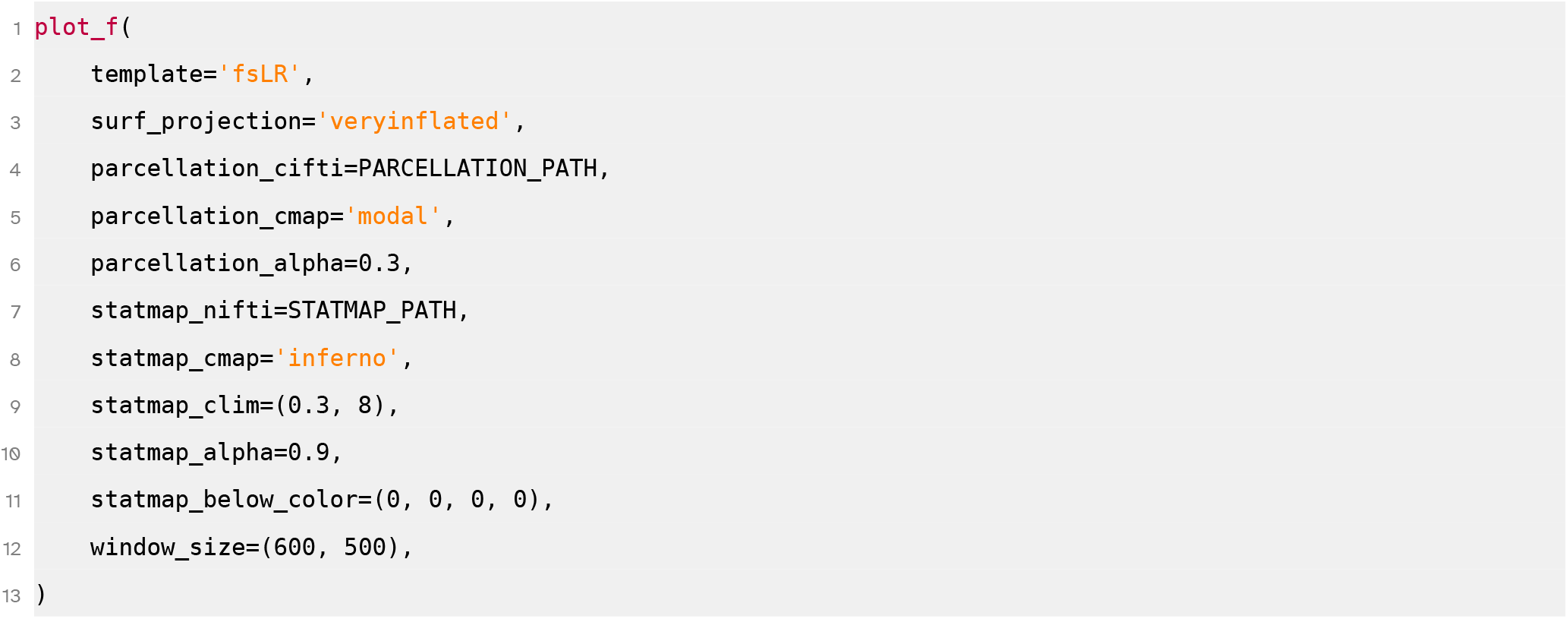

### 2.3 Output primitives

Output primitives determine the type of artefact that a visualisation protocol creates to represent a scene. They broadly fall into two categories: interactive and static.

#### Interactive outputs

Interactive scene representations include display windows and HTML files; they allow manipulation of camera angle and proximity. Display windows are created when invoking a visualisation protocol into which the `plot_to_display` primitive is composed. Display windows are fundamentally ephemeral and non-portable; they can be produced only if the original data are present. `hyve` can also create interactive scene representations that are persistent and portable, in the form of HTML files, using the `plot_to_html` primitive (Jourdain et al., 2023). This feature can be used to share interactive representations of scientific results.

#### Static outputs

Interactive scene representations are typically not compatible with legacy media such as print or static electronic publications. Static representations additionally carry the (dis)advantage of selectively focusing on salient features of a scene. A static visualiser is composed using the `plot_to_image` primitive. Static outputs are characterised by a fixed view angle; any visualisation protocol that contains a static output primitive will therefore acquire a `views` argument. `views` should be a sequence of views, each of which can be either a PyVista camera view specification or a string representing a direction along an anatomical axis, like `‘dorsal’`, `‘ventral’`, `‘anterior’`, or `‘posterior’`. If no `views` argument is explicitly supplied, 6 orthographic views are captured for each scene. `hyve` also contains “autocam” primitives that implement heuristics for automatically selecting one or more camera angles on a scene (Figure 5, detailed in Section 3.3.1).

**Figure 5:**
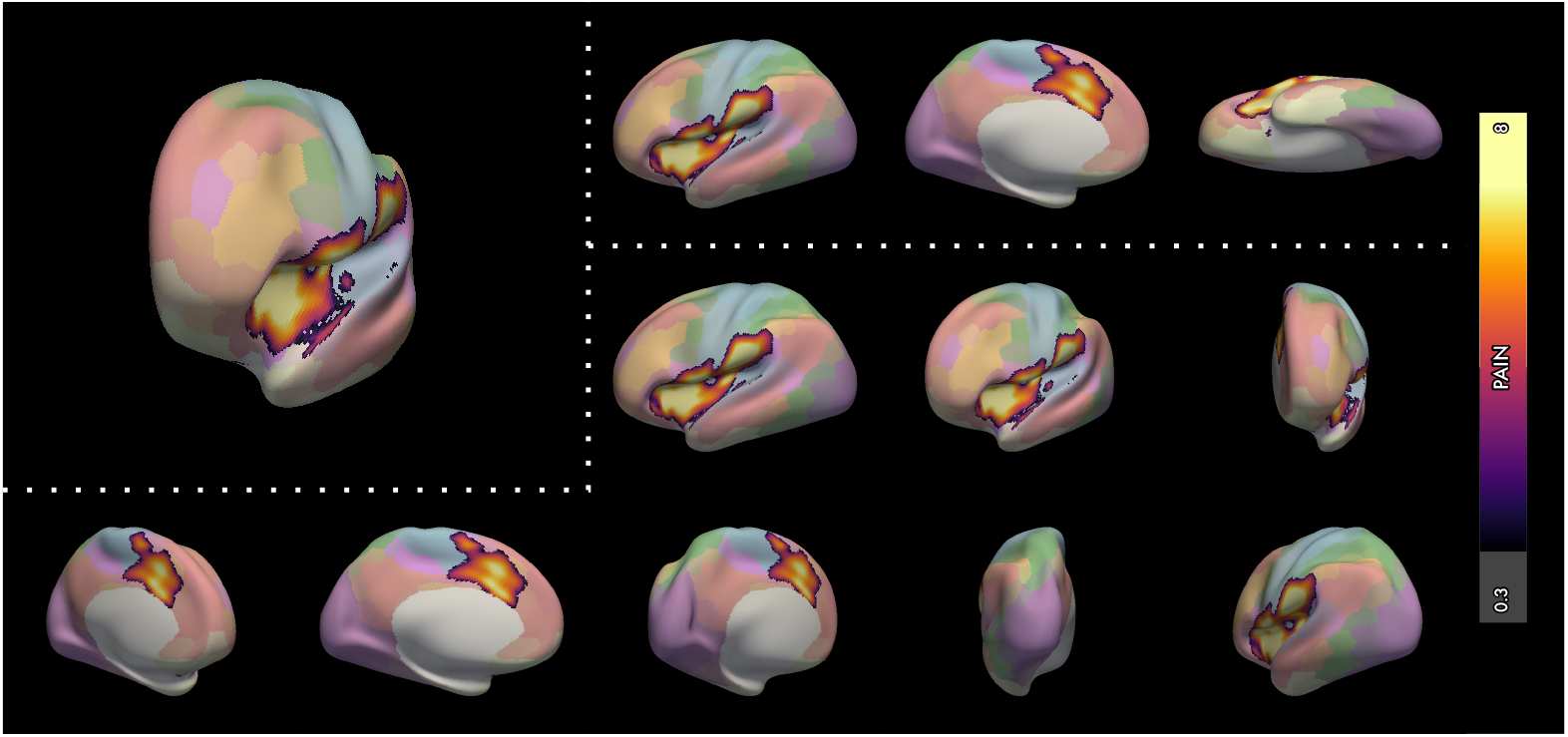
Examples of views selected by `hyve` autocam primitives. *Top left*, the `scalar_focus_camera` identifies the centroid of the meta-analytic mass and focuses the camera on it. *Top*, the `closest_ortho_camera` identifies three orthographic views closest to the peak of the scalars. *Bottom*, the `planar_sweep_camera` captures eight views through a turntable sweep of the scene. The number of view angles to capture is a parameter of the autocam primitive.

Static scene representations can be saved as individual snapshot files using the `save_snapshots` output primitive. `save_snapshots`, along with other output primitives that create persistent scene representations (like `plot_to_html` and `save_figure`), adds a `fname_spec` parameter. `fname_spec` is a Python string, optionally with brace-circumfixed format fields (e.g., `‘scalars-{surfscalars}_hemisphere-{hemisphere}_view-{view}’`). Each file that is saved has its format fields automatically populated from scene and view metadata (detailed in Section 3.1.2).

### 2.4 Figure builder

The core `hyve` plotting loop returns a sequence of scene representations together with a metadata dictionary for each scene. This scene-plus-metadata design undergirds a powerful, semantic *figure builder* API, which can be used to intuitively compose static scenes and other graphical elements into editable, publication-quality SVG figures. The figure builder comprises two components: a layout “algebra” and an annotation system. The end user interacts with the figure builder by (1) using the “algebraic” operations to specify a layout as a transformation of `hyve.Cell`s and (2) annotating the `Cell`s of this layout using tags present in the plot metadata. There is also an option to inject exogenous graphical elements, such as existing SVGs or functions that create `pyplot` figures.

The figure builder is invoked by any protocol containing a corresponding primitive—either `save_figure` or the convenience wrapper `save_grid`:

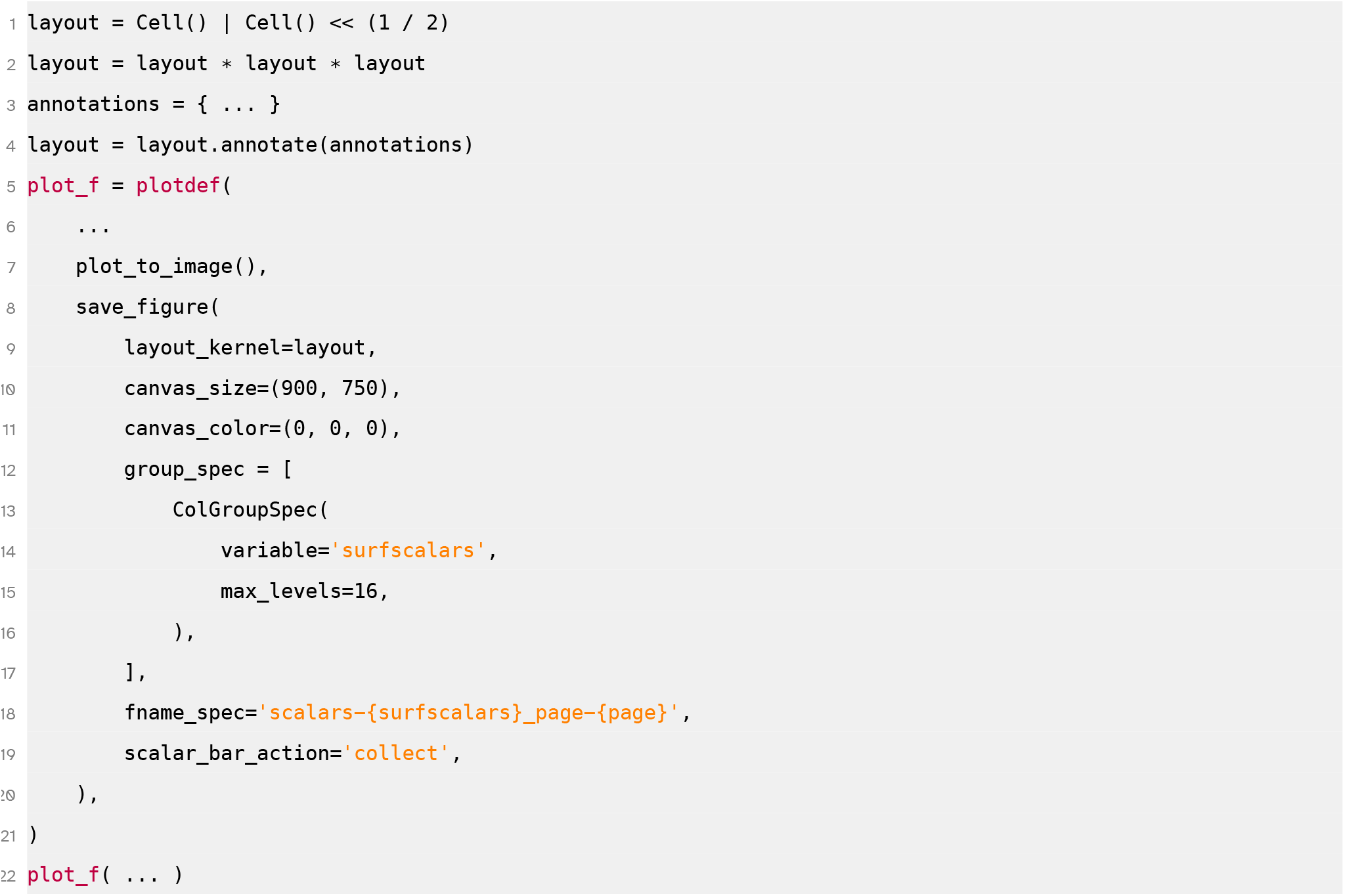

Parameterising a layout is simplified with a limited set of 6 noncommutative infix operations (the layout “algebra”, demonstrated by example in Figure 6 and detailed more formally in Section 3.4.2). The desired layout is assigned to the `layout_kernel` parameter, and further customisations are available through group capture specifications, discussed below.

**Figure 6:**
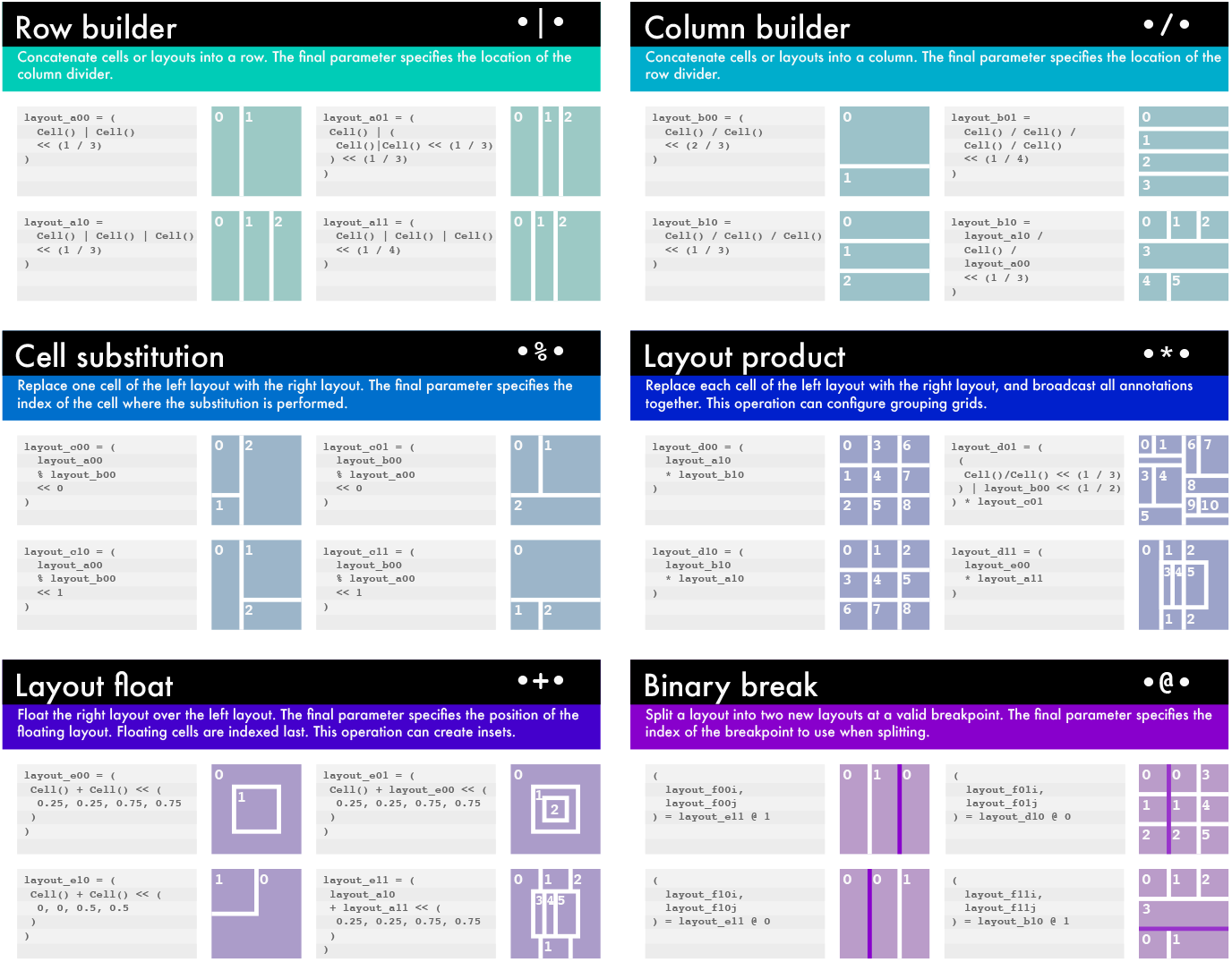
Overview of layout operations with examples. Figure layouts can be constructed from cells using six noncommutative infix operations. Further details are available in Section 3.4.2.

#### 2.4.1 Annotations

By default, the figure builder assigns each scene or graphical element into the first unas-signed cell in a layout. When designing figure layouts, however, we typically have in mind a particular purpose for each cell. Cell annotation reconfigures the procedure by which the figure builder primitive assigns scenes to cells so that it instead searches for matches between cell annotations and the semantic information in scene metadata (Figure 7). The cells of any layout can be annotated using a dictionary that specifies the desired contents of each cell:

**Figure 7:**
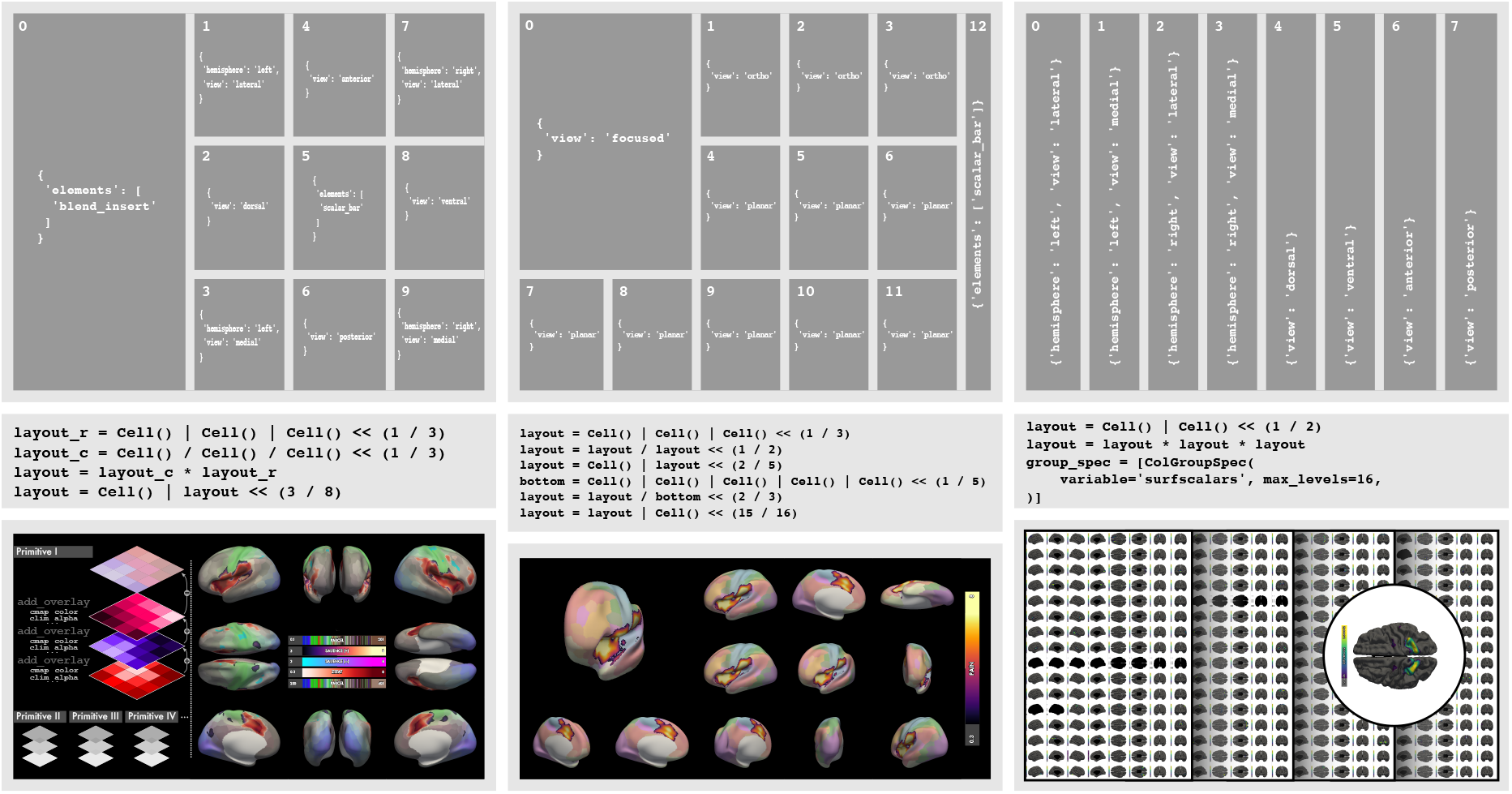
A layout can be annotated to semantically orchestrate figure construction. *Left* and *centre*, annotation (*top*) and configuration (*middle*) of figures 4 and 5 from this manuscript. Note that the figure at left contains an exogenous SVG element called `blend_insert`. *Right*, annotation and configuration of a multi-page figure using a group capturing specification. The figure displays eight orthographic views of 64 DiFuMo components (Dadi et al., 2020). The layout kernel (*top*) determines the configuration of each row, and the `ColGroupSpec` then multiplies the kernel over a column whose entries capture each surface scalar dataset in DiFuMo in a separate row. The resulting figure is spread over four pages, with 16 components plotted per page.

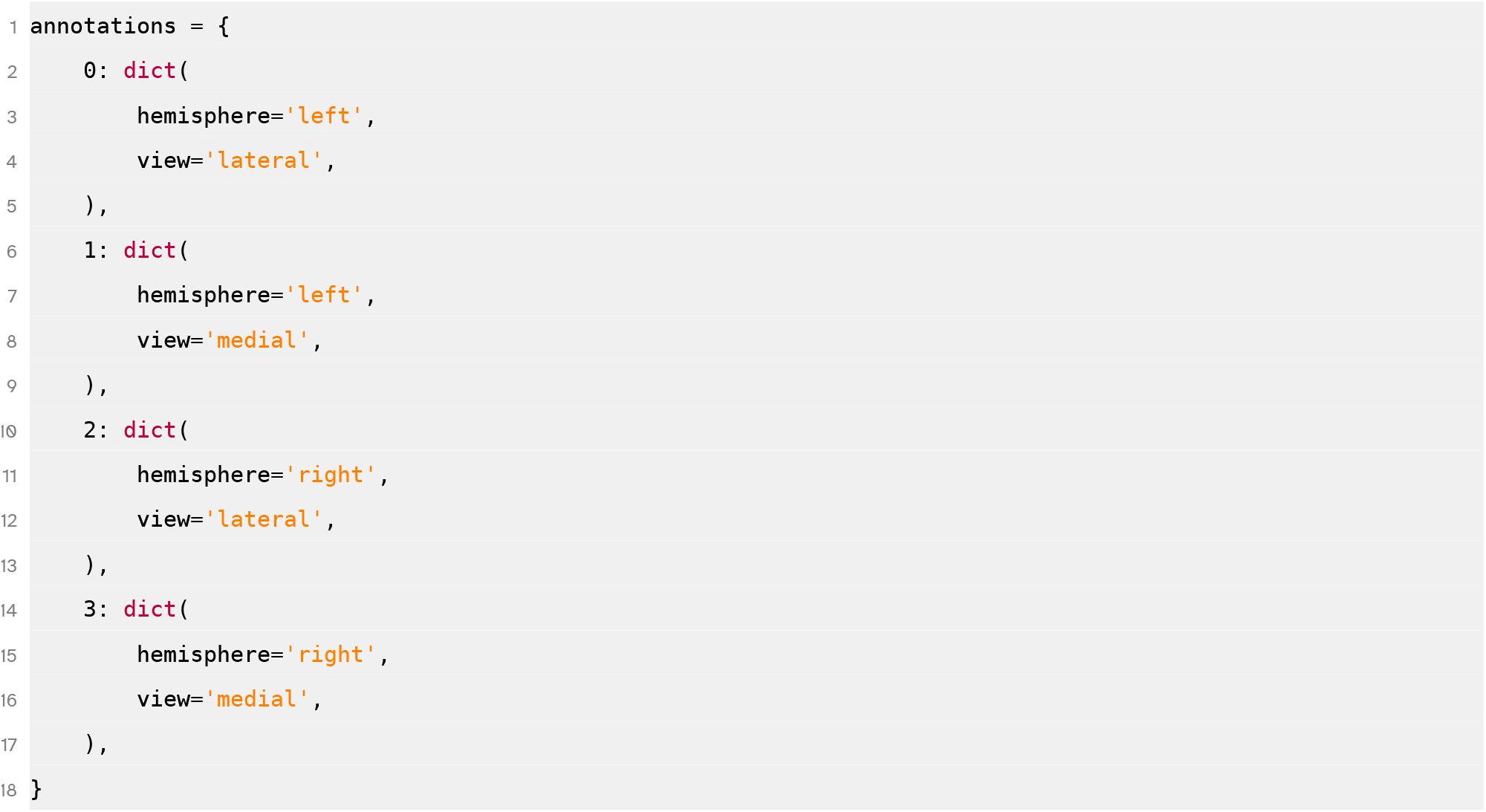

Occasionally a user might intend to create a figure that replicates a layout across multiple datasets, for instance components of a fMRI signal decomposition. `hyve` includes group capture specifications for this purpose. Group capture specifications capture all values of a metadata variable and replicate the layout kernel for each value (Figure 7, *right*). Group capture specifications can replicate layout kernels over a column (`group_spec=[ColGroupSpec(…)]`), a row (`group_spec=[RowGroupSpec(…)]`), or general grid (`group_spec=[GroupSpec(…)]`), and multiple captures can be sequentially applied for serial replication. Setting the `max_levels` parameter can limit the total size of a single figure, splitting excess panels over multiple pages.

#### 2.4.2 Exogenous elements

`hyve` includes input primitives that enable the figure builder to use graphical elements that are not returned as artefacts of the core plotting routine. This functionality supports, for instance, construction of SVG elements like text boxes (`text_element` primitive), embedding of existing SVG files into layout cells (`svg_element` primitive), and injection of PyPlot figures, such as those produced by `matplotlib` (Hunter, 2007), `seaborn` (Waskom, 2021), or `nilearn` (Abraham et al., 2014) (`pyplot_element` primitive).

### 2.5 Tutorial

A tutorial for new `hyve` users is available on GitHub at

https://github.com/hypercoil/notebooks/blob/main/nb/hyve/hyve-constructive.ipynb

## 3 Software implementation reference

### 3.1 Design and dependencies

The `hyve` API is built around the function `hyve.plotdef`, which transforms an abstract visualisation loop into a concrete protocol for creating a specific visualisation artefact. `hyve`’s base plotter function contains fewer than 100 lines of code and performs only abstract operations: (i) initialising a scene, (ii) transforming geometric datasets, (iii) interpreting geometric datasets as scene “actors” using *geometric primitives*, and (iv) optionally applying *postprocessor* callables to create a serialised representation of the scene, which is returned alongside instructions for building scene-dependent vector graphic elements like scalar bars. These abstract operations are made concrete when the base plotter is composed with geometric primitives. Each geometric primitive specifies an instruction set for the base plotter to interpret datasets structured by a particular class of geometry: for instance, a surface, a point cloud, or a network. The core visualisation loop that each visualisation protocol calls is already composed with all available geometric primitives; the separation of abstract routines from geometry-specific implementations is a design choice to improve the software’s future extensibility to new geometries.

#### 3.1.1 Scope, limitations, and relationship to existing work

The compositional paradigm aligns with `hyve`’s intended design as an extensible *visualisation engine*. The role of a visualisation engine is to expose an API that facilitates defining new high-level visualisation functions for specific purposes. In `hyve`, these definitions are instantiated using the `plotdef` function, which takes as positional arguments a sequence containing any number of parameterised *transforms*. For simplicity, we have used the term “primitive” loosely throughout Section 2 to refer to these transforms, but there is an important distinction—a transform is an instruction set for composing a particular primitive into the visualisation protocol. Chaining transforms together in `plotdef` constitutes a flexible and modular framework for composing new visualisation protocols that utilise the `hyve` core plotting loop. `hyve.plotdef` returns a new function, whose documentation (shown when calling `help(f)` on function `f`) is automatically constructed from its signature.

A clear drawback of this approach is a steeper learning curve compared with visualisation libraries that include a preconfigured subset of plotting routines out of the box, or compared with GUI-centred systems (but see also Chopra et al., 2023). However, the level of abstraction is intermediate between such high-level libraries (e.g., Fanton and Thompson, 2023; Gale et al., 2021) and more general frameworks like PyVista (Sullivan and Kaszynski, 2019) or MayaVi (Ramachandran and Varoquaux, 2011). Another notable example of a visualisation engine for brain data is VisBrain (Combrisson et al., 2019), which adopts an object-centred framework in contrast with `hyve`’s functional approach.

Aside from the possibility of composing invalid protocols, the flexibility offered by the functional approach comes with some notable constraints. When a new protocol `plot_f` is composed via `hyve.plotdef`, `plot_f` accepts only keyword arguments, and as a closure it might not behave reliably when serialised.

#### 3.1.2 The core visualisation (and metadata) loop

`hyve` wraps calls to its base plotter in a loop that automatically maps sets of argument assignments over separate plotter calls, thereby enabling the sequential creation of multiple scenes. Within this loop, each plotter call is also paired with a corresponding call to a metadata dictionary builder. Like the base plotter, the metadata builder is a minimal, abstract function that is built up through composition with geometric primitives. The composite metadata builder returns a dictionary containing semantic information about each call’s argument assignments, which communicates the expected contents of each scene.

To map over plotter calls, each argument to the core loop can be assigned a sequence of values instead of a single value. The core loop of `hyve` additionally accepts a `map_spec` argument: this *mapping specification* instructs the core loop how it should select and combine values from sequential arguments for each call made to the base plotter (Figure 8). It uses a simple syntax inspired by the workflow library `pydra` (Jarecka et al., 2020): the names of parameters whose values are sequences to be mapped over are enumerated in a data structure that we define recursively here. The base case of this data structure is a single variable name. Then, any Python list or Python tuple of mapping specifications is a valid mapping specification. For specifications that are in the same *list*, the core loop performs an *outer map*, creating every possible combination of assignments to those specifications. For specifications that are in the same *tuple*, the core loop performs a *scalar map* by associating each assignment to one specification with a corresponding assignment to each other specification. For each inferred assignment combination, the core loop makes a call to the base plotter and the metadata builder (Figure 8).

**Figure 8:**
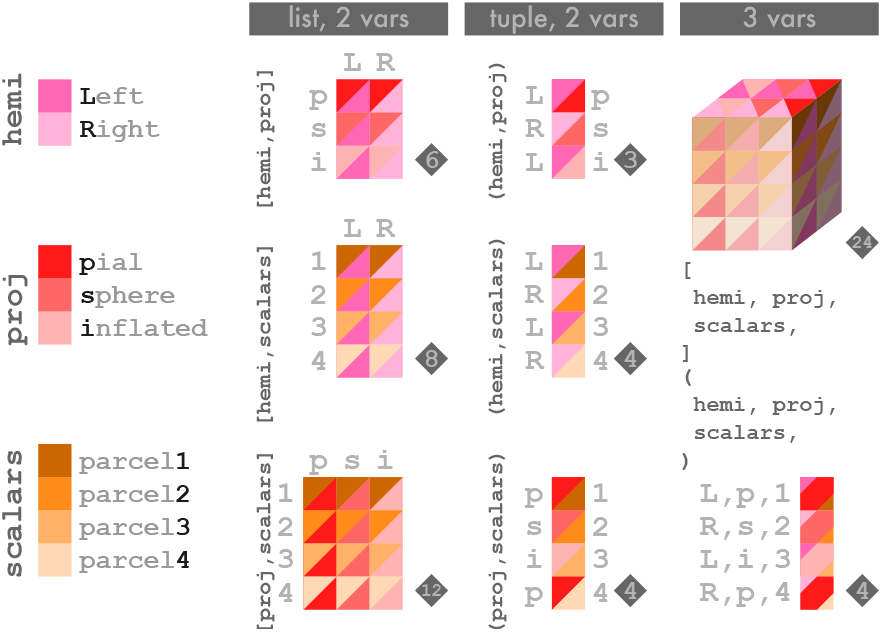
Schematic illustrating the argument mapping protocol of the core visualisation loop. When the core loop receives an argument whose value is a sequence, it maps the values in that sequence across multiple calls to the base plotter. If multiple arguments are supplied as sequences, the core loop must decide whether particular argument values should always be paired or whether all possible combinations of assignments should be submitted in separate base plotter calls. The user can control this behaviour by designating a *map specification* (`map_spec`), which can be either a tuple or a list that contains either argument names or nested map specifications. A list indicates that the full “outer” set of combinations should be generated, while a tuple indicates that values sharing the same index should be paired. The illustration shows how different map specifications group argument values together and indicates the total number of base plotter calls in each scenario.

#### 3.1.3 Installation and dependencies

`hyve` is installable via `pip` from PyPI for Python 3.10+. We have opted to use PyVista (Sullivan and Kaszynski, 2019) as the main rendering backend. PyVista is a modern library that exposes a “Pythonic” interface to VTK, a workhorse scientific 3D graphics system with support for GPU acceleration (Schroeder et al., 2006). PyVista is also used as the visualisation backend for MNE, the gold standard Python library in EEG and MEG signal processing (Gramfort, 2013).

Installation of `hyve` can be challenging on some systems due to its dependency on VTK. For those systems, `matplotlib`-based (Hunter, 2007) visualisation libraries like `netplotbrain` (Fanton and Thompson, 2023), `brainplotlib` (Ma et al., 2022), or `nilearn`’s plotting annex (Abraham et al., 2014) offer a suitable alternative. `hyve`’s Python dependencies can be separated neatly into four categories:

- First, the core scientific Python suite—`numpy`, `scipy`, `pandas`, and `matplotlib`;
- Second, visualisation interfaces—`pyvista`, `trame` and `trame-vtk` (for creating interactive HTMLs; Jourdain et al., 2023), and `svg.py` (for creating editable SVG figures);
- Third, specialised neuroimaging libraries—`nibabel` (for reading and writing common neuroimaging data formats), `templateflow` (for accessing template geometries and atlases; Ciric, Thompson, et al., 2022), and `lytemaps` (a `pooch`-based fork of `neuromaps`, for resampling data and accessing certain maps not available in `templateflow`; Markello et al., 2022; Uieda et al., 2020); and
- Finally, the ultra-lightweight `conveyant` library, another component of the `hypercoil` project that contains lower-level abstractions for compositional functional programming.

We took specific care not to require omnibus analysis libraries such as `nilearn` as dependencies given the comparatively specific focus of `hyve` as a visualisation engine.

### 3.2 Geometric and input primitives

Although the end user does not directly interact with geometric primitives, they perform a critical function in the `hyve` library by instructing the base plotter and base metadata builder how to interpret datasets structured by different types of geometries. For the initial release of `hyve`, we focused on exposing geometric primitives that would cover some of the most common use cases in neuroimaging. However, the compositional design of the base plotter means that extending `hyve` to support additional primitives is relatively straightforward. In particular, PyVista additionally supports *inter alia* surface contours, vector fields, volumetric blocks with cutaways and slices, surface mesh reconstructions of volumetric regions of interest, streamlines, and numerous relevant geometries that could be encapsulated in future primitives (Sullivan and Kaszynski, 2019).

#### 3.2.1 Surfaces and point clouds

Surface datasets (Figure 2) are structured by mesh elements—either faces or vertices— on an instance of the `ProjectedPolyData` class, typically one per cortical hemisphere. `ProjectedPolyData` is a subclass of PyVista’s `PolyData` with additional support for storing multiple surface *projections*. The availability of surface projections varies from template to template, but common types include pial surfaces that follow the contours of cortical gyrification, inflated surfaces that better expose sulcal mesh elements, spherical surfaces that simplify computations while preserving homotopy, and exploded flat projections that embed all mesh elements in a two-dimensional plane. Point cloud scalars (Figure 3, *top*) are internally represented as unstructured point sets.

#### 3.2.2 Brain networks

Because there is little consensus about data formats for representing and storing brain network data, `hyve` instead accepts either raw `numpy` arrays or `pandas.DataFrame`s. The `build_network` input primitive, which constructs `hyve`’s internal container for network geometries, requires each network to be represented as two `DataFrame`s and one `numpy` array. The `numpy` array specifies the coordinates of each node, while the two `DataFrame`s store node-level and edge-level data. The coordinate array is either set explicitly or obtained from centroids of a surface scalar parcellation dataset using the input primitive `node_coor_from_parcels`. The indices of the node-level `DataFrame` are integers corresponding to the respective rows of the coordinate array. To allow consistency with label numbers in common parcellations, 1-indexing is used instead of 0-indexing. The edgelevel `DataFrame` has two index columns: one named `src` and another named `dst`, each containing the index of one of the two nodes that the edge connects. Each `DataFrame` can also have any number of additional columns. The names of these columns become available as values to aesthetic parameters like `node_alpha` or `edge_radius`.

The `add_node_variable` and `add_edge_variable` primitives, by comparison, are used to build `DataFrame`s from `numpy` arrays containing data. For `add_node_variable`, the input data array is a single vector; for `add_edge_variable`, it is a square adjacency matrix. These primitives additionally implement filtering operations like thresholding, value substitution, decomposing a signed variable into a magnitude variable and a sign variable, selecting edges that connect to a subset of nodes or nodes on which edges are incident, and emitting a node degree dataset from an input adjacency matrix.

Brain network scenes containing tens or hundreds of thousands of edges are challenging to render in a reasonable time. Internally, `hyve` leverages PyVista’s glyph functionality to dramatically accelerate rendering of networks. This means that nodes and edges are plotted as point clouds represented by spheres and cylinders, respectively, and those spheres and cylinders are oriented appropriately and scaled according to specified data attributes.

#### 3.2.3 Overlays and blending

Through the use of overlays and blending (Figure 4), it is possible to structure multiple datasets using a single geometry in a single scene. `hyve`’s `add_overlay` input primitives disambiguate between (a) the default behaviour of mapping over multiple input datasets (see Section 3.1.2) and (b) an alternative behaviour of overlaying multiple input datasets. Because adding overlays also adds aesthetic arguments to the plotting protocol, it is clearly necessary to designate a unique name for each overlay in order to prevent parameter collisions.

`hyve` constructs a representation of a geometric dataset by progressively transforming an RGBA tensor defined over the elements of the geometry (e.g., vertices, faces, points, or glyphs). First, it constructs a base layer from generic parameter values like `surf_scalars_cmap`. The base layer is constructed by using the assigned colour parameters to map scalar values into RGBA space. Next, for each successive overlay, a similar RGBA tensor is constructed from the corresponding parameter values (e.g., `parcellation_cmap`, `statmap_cmap`). As each tensor is constructed, the source-over compositing heuristic is applied with premultiplication of each layer’s alpha channel (Ciechanowski, 2019), ultimately producing a single RGBA tensor representation containing the blended values of all overlays. This final composite representation is then structured by the geometric primitive.

When multiple overlays are associated with a single structuring surface, `hyve` requires that the corresponding datasets be defined over a common kind of mesh element—for example, either vertices or faces, not some combination of both. `hyve` provides `vertex_to_face`, a primitive for resampling vertex-valued data onto face polygons to facilitate this.

### 3.3 Output primitives

An interactive scene representation is characterised by a capacity to manipulate the view angle. Additional interactivity (e.g., creating cutaways or retrieving information from a mesh element) could potentially be provided by widget primitives in future extensions of `hyve`, but these are not yet implemented. Interactive visualisation modes currently implemented in `hyve` are display windows and HTML files. In the terminal, the `plot_to_display` primitive will create a window containing a live rendering of the scene. In a Jupyter notebook, this will create an inline `trame` visualisation panel. The `plot_to_html` primitive uses PyVista and `trame` (Jourdain et al., 2023) to create interactive HTML plots.

By contrast, scenes are statically represented as PNG images. `plot_to_image` only stages PNG representations in an in-memory buffer. This is so they can be either saved individually or laid out as panels in a figure (using either an existing layout system like `matplotlib.gridspec` or `hyve`’s native semantic SVG figure builder API, described in the next section).

#### Autocam primitives

“Autocam” (automatic camera) primitives use a heuristic or rule to select a view or views on the scene (Figure 5). Three autocam primitives, currently designed for use with surface geometries, are included with the initial release of `hyve`:

- The `scalar_focus_camera` primitive determines either the peak or the centroid of a displayed surface scalar dataset. It then places the camera along a vector drawn from the physical centre of the surface through the peak or centroid coordinates.
- The `closest_ortho_camera` computes the distance from the peak or centroid to surface “poles”—extremal coordinates along each spatial axis. The distance computation is performed on a spherical projection if one is available for the current dataset. It then places a camera pointing toward each of the closest *n* poles of the surface, where *n* is a primitive parameter.
- The `planar_sweep_camera` (turntable) requires an initial camera position vector, a normal vector that is orthogonal to the initial vector, and an integer *n*. Starting from the initial vector, it computes *n* views toward the scene centre, all from the plane that is parameterised by the normal vector.
- An additional `auto_camera` primitive is included as a convenience function that allows for easy combination of autocams.

### 3.4 Figure builder

When a figure builder output primitive is composed into the visualiser loop, `hyve` will automatically search the metadata returned by the core plotting routine to bind particular scenes and graphical elements to matching layout cells, according to the annotations present.

#### 3.4.1 Layout data structure

Figure layouts in `hyve` are represented as binary trees (Figure 9). Each layout has two children that are also either layouts or leaf nodes. Every leaf node is a single instance of the `Cell` class. Every non-leaf node is associated with two parameters: a *split location* and a *split orientation*. The figure builder uses split locations and orientations to partition a rectangular canvas into the cell layout parameterised by the tree schema; each node of the tree thus corresponds to a rectangular patch of the canvas. A vertical split orientation divides a canvas patch vertically such that its children form a row, while a horizontal split orientation divides a patch horizontally such that its children form a column. The split location is a fraction between 0 and 1 that specifies the relative sizes of the child patches. Layouts are defined abstractly, without a fixed size. Calling the `partition` method of a layout with two integer arguments (*x, y*) returns a layout whose cells additionally have their `loc` and `dim` properties populated with the cells’ offsets and sizes in pixels for a layout of that total dimension. The binary tree representation of a layout induces a natural ordering over the cells in that layout (Figure 9).

**Figure 9:**
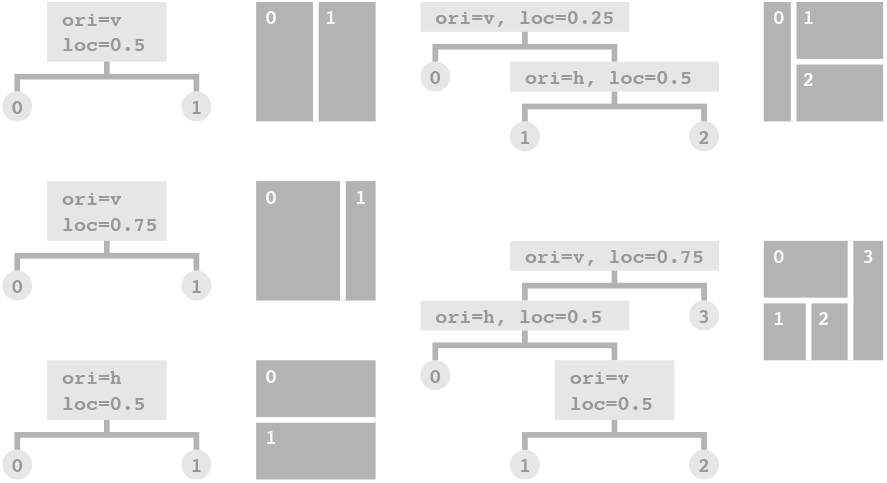
Duality of binary trees and canvas layouts. For five binary trees whose non-leaf nodes have split orientation (`ori`) and location (`loc`) properties, the corresponding canvas layout is shown, together with the cell ordering induced by the binary tree.

#### 3.4.2 Layout algebra

The “algebra” used to configure layouts comprises six elementary, noncommutative infix operations. Four of these operations are additionally parameterised by an additional numeric value that is offset by the left-shift operator (<<).

- *Row-builder* (|). The | operation is used to concatenate layouts (including cells) into a row. A row concatenation expression takes the form L_0[ | L_i]+ << s, where the bracketed expression can be repeated indefinitely, the *L*_*i*_s can be any layout, and *s* ∈ (0, 1). When *n* total layouts are concatenated, 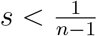 must be true. The expression returns a new layout whose descendants include all of the concatenated layouts, and in which the split locations introduced by the concatenation are positioned such that all concatenated layouts except for the last take up a fraction *s* of the total horizontal dimension of the new layout; the last layout takes up whatever fraction is remaining.
- *Column-builder* (**/**). The / operation functions much the same as the row-builder, except that it concatenates layouts vertically into a column.
- *Cell substitution* (**%**). The % operation is used to replace a single cell of one layout with the entire structure of a second layout. A cell substitution expression takes the form L_0 % L_1 << i, where integer *i* ∈ [0, |*L*_0_|) for |*L*_0_| the total number of cells (leaf nodes) in layout *L*_0_. The expression returns a new layout whose structure matches *L*_0_, except with the *i*th cell replaced with the structure of layout *L*_1_.
- *Layout product* (**\***). The * operation replaces each cell of one layout with the entire structure of a second layout. A layout product operation takes no extra parameters and has the form L_0 * L_1. This operation can be used to quickly create grids, by taking the product of a row layout and a column layout. Noncommutativity implies that row- or column-major indexing follows from the order of the operands. The layout product has an additional behaviour when applied to annotated layouts: each cell in the product layout inherits annotations from its analogue in both operands. This enables the use of the product as a grouping function, for instance to group figure panels according to a metadata variable like brain hemisphere or view angle.
- *Float* (**+**). The + operation “floats” one layout over another, creating an inset panel. A float expression takes the form L_0 + L_1 << (x_0, y_0, x_1, y_1)), where *x*_0_ ∈ [0, 1) and *x*_1_ *> x*_0_, *x*_1_ ∈ (0, 1] set the coordinates of the left and right boundaries of the floating layout *L*_1_ relative to the fixed layout *L*_0_. Similarly, *y*_0_ ∈ [0, 1) and *y*_1_ *> y*_0_, *y*_1_ ∈ (0, 1] set the coordinates of the top and bottom boundaries. A floating layout cannot contain another floating layout; when a layout *L*_1_ containing a floating layout *L*_2_ is floated on *L*_0_, the “inner” floating layout *L*_2_ is also “refloated” on the same fixed layout *L*_0_ by changing the reference frame of its float coordinates. Because floating layouts are not intrinsically part of the binary tree structure of the fixed layout, their cells are indexed after all cells in the fixed layout. When multiple floating layouts are present, they are indexed in the order that they were floated.
- *Binary break* (**@**). The @ operation breaks a single input layout into two new layouts. A binary break expression takes the form L_0 @ j, where *j* indexes a valid breakpoint of the layout. Valid breakpoints are defined recursively following the tree structure of the layout: as a base case, the root node of the tree is always a valid breakpoint. Recursively, for any node that is a valid breakpoint, immediate child nodes whose split orientations match that node’s split orientation are valid breakpoints. This recursive definition has an intuitive realisation in canvas layouts: it means that valid breakpoints are any split lines that run across the entirety of the canvas. Breakpoints, like cells, are indexed according to the order induced by the binary tree representation. The binary break operation can be useful for dividing a large and complex figure into multiple pages. Note that binary break operations will automatically drop any floating layouts that straddle the break. Also note that binary breaks should only be applied to deliberately constructed layouts, because split locations are defined relative to the immediate parent and undesired behaviour will result in other cases.

#### 3.4.3 Annotations

Valid annotation terms are identical to the key-value pairs in scene metadata. These key-value pairs differ based on the geometric primitives present in a scene and the arguments supplied to each call—for instance, when the core plotter is mapped over different values of the `hemisphere` argument or over different `surfscalars`, the specific argument value assigned to each core plotter call is also entered into a corresponding metadata dictionary, thereby semantically associating each scene representation artefact produced by the core plotter with a set of metadata.

The figure builder uses a greedy heuristic to assign scene representations to layout cells according to metadata and annotations present. For each scene representation, annotations filter the availability of cells as candidates for placement of that scene representation: if a cell contains an annotation that conflicts with the scene’s metadata, that cell is then ruled out as a candidate for placement of that scene. The assignment difficulty for each cell is defined as the total number of annotations present in that cell: the more annotations are present, the greater the number of successful matches required to place a scene there. After determining candidates and ranking them by assignment difficulty, the figure builder will place each scene in the first candidate cell with the highest assignment difficulty. After all scene representations that can be assigned to cells in this way have been assigned, the remaining representations can optionally be forced into the remaining cells.

## 4 Discussion

We developed `hyve` as a free and open-source visualisation engine for neuroimaging in Python. In comparison with existing software solutions, `hyve`’s function-oriented design centres on compositional construction of new visualisation protocols from atomic primitives. At `hyve`’s core is an abstract visualisation loop that automatically maps plotter calls over multiple inputs and returns scene metadata for each call. The abstract core loop is transformed into concrete protocols for visualisation through `hyve`’s primary user interface, `plotdef`, which composes primitives that impart specific functionalities into the loop. The visualisation functions produced by `plotdef` are reusable and automatically documented. `hyve`’s compositional paradigm underpins a modular and extensible library that supports visualisation of common classes of geometrically structured data, including surface representations of the cerebral cortex, point-cloud representations of image volumes, and glyph representations of brain networks.

The compositional design of `hyve` further enables configuration of new visualisation functions that handle common input data formats and research objectives. In addition to ingress of data in standard neuroimaging data formats like NIfTI and GIfTI, the software’s input primitives also support common use cases like viewing parcellated data or retrieving surface geometries representing a standard space from a cloud-based archive. `hyve` also includes a set of filters for brain network datasets formatted as adjacency matrices.

`plotdef` parameterises functions that produce visualisation artefacts that can be static or interactive, persistent or ephemeral. Interactive visualisations allow a client to manipulate a scene and include display windows as well as HTML files that are persistent and portable. Static visualisations, by contrast, support configuration of fixed views on scenes. `hyve` also includes a figure builder API that combines a flexible layout algebra with a simple annotation system to support semantic composition of editable multi-panel figures.

Because a visualisation engine is highly configurable, and because the functional paradigm might be unfamiliar to users who are better acquainted with imperative and object-oriented programming patterns, we also provide a tutorial for `hyve` in the form of a Jupyter notebook. This tutorial takes a constructive approach, stepping users through the process of building a new visualisation protocol by adding and configuring individual primitives. It is available on GitHub at https://github.com/hypercoil/notebooks/blob/main/nb/hyve/hyve-constructive.ipynb and additionally requires the installation of the `hyve-examples` package from PyPI.

### 4.1 Limitations and extension

`hyve`’s limitations are of two broad categories: (i) absent or limited support for certain geometries and configurations and (ii) a relatively steep learning curve. Support for geometries such as slices, streamlines, and reconstructions is present in PyVista, the visualisation library that underlies `hyve`, and future support is therefore a matter of exposing these to the visualisation loop through new geometric primitives. Similarly, although support for different templates and other features is limited at this time, extending the available options is a matter of quantitative rather than qualitative change. The steeper learning curve is a necessary consequence of configurability; it follows from the role of `hyve` as a neuroimaging visualisation engine that sits at an abstraction level intermediate between general frameworks like PyVista and preconfigured plots. The demand for higher-level turnkey interfaces could be addressed by future development of a `hyve`-based library that implements a gallery of useful plot configurations.

## Ethics statement

All data used in this work were retrieved from public repositories; additional information regarding the collection of these data and relevant regulations can be found in cited work. No new data were collected specifically for the development of the software library described here.

## Data and code availability

The most recent stable release of `hyve` is available for installation from PyPI using `pip`. The source code is available on GitHub at https://github.com/hypercoil/hyve/; the latest development version can also be installed from the GitHub repository.

## Author contributions

R.C. developed the software library and wrote the initial draft of the manuscript. A.X. designed additional test cases, provided feedback to guide development, and revised the manuscript. R.A.P. funded development, provided feedback to guide development, and revised the manuscript.

## Supplemental material

An introductory tutorial can be downloaded from https://github.com/hypercoil/notebooks/blob/main/nb/hyve/hyve-constructive.ipynb.

